# Optimising the viscoelastic properties of hyaluronic acid hydrogels through colloidal particle interactions: a response surface methodology approach

**DOI:** 10.1101/2024.08.12.607572

**Authors:** Giriprasath Ramanathan, Masroora Hassan, Yury Rochev

## Abstract

Enhancing the viscoelastic characteristics of hydrogel systems through strategic colloidal particle interactions is paramount for their functionality in rectal gel applications. This investigation delves into the synergistic interactions between cationic nanoparticles (CNPs), anionic nanoparticles (ANPs), and composite nanoparticles (NPs) within a hyaluronic acid (HA) hydrogel matrix, employing response surface methodology (RSM) for optimisation. Critical parameters, namely the volume fraction of NPs and oscillatory amplitude, were meticulously calibrated to achieve optimal complex viscosity, as determined by advanced rheometric analysis. The findings reveal substantial effects of CNPs, ANPs, and mixed NPs on the viscoelasticity of the HA hydrogel, with complex viscosity measurements of 490.82 ± 10.57, 214.70 ± 8.96, and 328.46 ± 6.67 mPa·s, respectively. The hydrogel system with mixed NPs exhibited a strong concordance with empirical data (R² = 0.9843), validating the predictive precision of the model. Morphological assessments uncovered a highly interconnected network within the HA gel-particle composite, characterised by both densely and sparsely packed porous architectures. This study presents a robust framework for modulating viscoelastic properties in colloidal particle-gel systems, providing pivotal insights for the development of advanced rectal gel formulations.

## 1. Introduction

The interaction between colloidal particles and hydrogels has garnered significant attention because of its potential applications in various fields, including nanocomposite polymers, rectal gel [1], 3D printing [2], and environmental engineering [3]. Hydrogels are solvent-inflated three-dimensional (3D) microstructural networks. The formation and properties of these hydrogels within polymer solutions depend on the characteristics of colloidal particles, such as size, surface charge, and physicochemical interactions [4]. The rheology of nanoparticles (NPs) within a 3D hydrogel network can display advanced transport properties, thixotropic behaviour, nonlinear mechanical properties, and micro- and nano-scale structural anisotropy. These properties significantly affect the material’s viscosity by balancing attractive and repulsive forces [5]. The hydrogel microstructure can be physical or chemical interactions between the polymer and NPs. These interactions can be attractive or repulsive, depending on various forces such as van der Waals, electrostatic, depletion, steric, hydrophobic, bonding, capillary, and magnetic forces. These forces are determined based on the DLVO (Derjaguin-Landau-Verwey-Overbeek) theoretical model [6]. The mechanical properties of colloidal gels formed by physical interactions can be distinctive because of their disordered microstructure [7] [8].

Ulcerative colitis (UC) is an inflammatory bowel disease (IBD) that causes chronic symptoms and leads to the erosion and damage of rectal, distal colonic, or entire colonic mucosa, submucosa, and crypt layers [9] [10]. The rectal administration of a colloidal suspension of nanoparticles [1], traditional plant extract [11], and drugs [12] in a hydrogel can enhance nanoparticle retention in the crypts, improving therapeutic efficacy [9] [13].

Hyaluronic acid (HA) is a hydrophilic polysaccharide characterised by carboxyl and hydroxy groups, capable of forming a 3D network gel with viscoelastic properties and high-water retention through intramolecular hydrogen bonding [14]. HA solutions are non-Newtonian fluids exhibiting shear thinning behaviour, where increasing shear rates disrupt intermolecular interactions, leading to reduced viscosity [15]. Commercially available rectal gel products, used in drug delivery systems, typically exhibit complex viscosities ranging from 145 to 300 mPa.s, ensuring drug stability and efficacy across various formulations [16] Previous studies have shown a decrease in gel viscosity when using chitosan/poly(vinyl alcohol) hydrogels with negatively charged NPs [4] and negatively charged silica NPs in acrylamide polymer systems[17]. Conversely, positively charged chitosan NPs in HA hydrogels significantly increased viscosity by enhancing interactions between hyaluronic acid chains, resulting in a stiffer and more rigid material [18].

This research explores the effects of cationic and anionic particles, as well as their synergistic combinations, on the viscosity of native HA gel using response surface methodology (RSM) with a Central Composite Design (CCD). To the best of our knowledge, the utilisation of RSM to investigate the interplay between nanoparticle surface charges and the HA gel system for achieving the desired complex viscosity has not been reported. This study investigates the interrelationship among process parameters by implementing CCD to understand HA gel-particle interactions and optimise the enema gel’s properties.

## 2. Experimental section

### 2.1. Materials

High molecular weight (HMW) Hyaluronic acid (HA) or sodium hyaluronate (Mw 1250-1500 kDa), purchased from Contipro, Czech Republic. PLGA (Resomer RG 504H, poly (D, L-lactide-co-glycolide), 50:50, Mw 38-54 kDa), Dichloromethane (DCM), Ethanol (ETOH), Polyvinyl alcohol (PVA) (Mw 13-23 kDa), Poly-L-Lysine (PLL) solution (Mw 150-300k Da, 0.1% w/v in H2O), and Phosphate buffer saline (PBS) (pH 7.4) were obtained from Merck. All other chemicals are from Merck unless specified otherwise.

### 2. 2. Synthesis of the HA functionalised nanoparticles

The HA-functionalized polymeric nanoparticle system was fabricated using PLGA as the core, followed by coatings with a primary layer of cationic polymer (PLL) and a secondary layer of anionic polymer (HMW-HA) **(Scheme 1(A))**. Briefly, PLGA (2% w/v) was dissolved in a DCM-ETOH (3:1) solvent mixture. This organic phase (1 mL) was added drop-wise to an aqueous phase containing 1% (w/v) PVA solution (10 mL) while being sonicated (3 min, pulse on/off: 8 s/2 s) in an ice bath. The organic solvents were evaporated by overnight stirring, followed by centrifugation at 10000 rpm for 10 minutes. For the primary coating, the PLGA NPs pellet was treated with a 0.1% (w/v) PLL solution. After 24 hours of stirring, the mixture was centrifuged (10000 rpm, 10 min). For the secondary coating, HA (0.2% w/v) was added to the PLL-coated NPs. After 6 hours, the PLGA-PLL-HA NPs were centrifuged (10000 rpm, 10 min), lyophilised, and stored [19]. Henceforth, the terms cationic nanoparticles (CNPs) and anionic nanoparticles (ANPs) refer to PLGA-PLL NPs and PLGA-PLL-HA NPs, respectively.

**Scheme 1.**
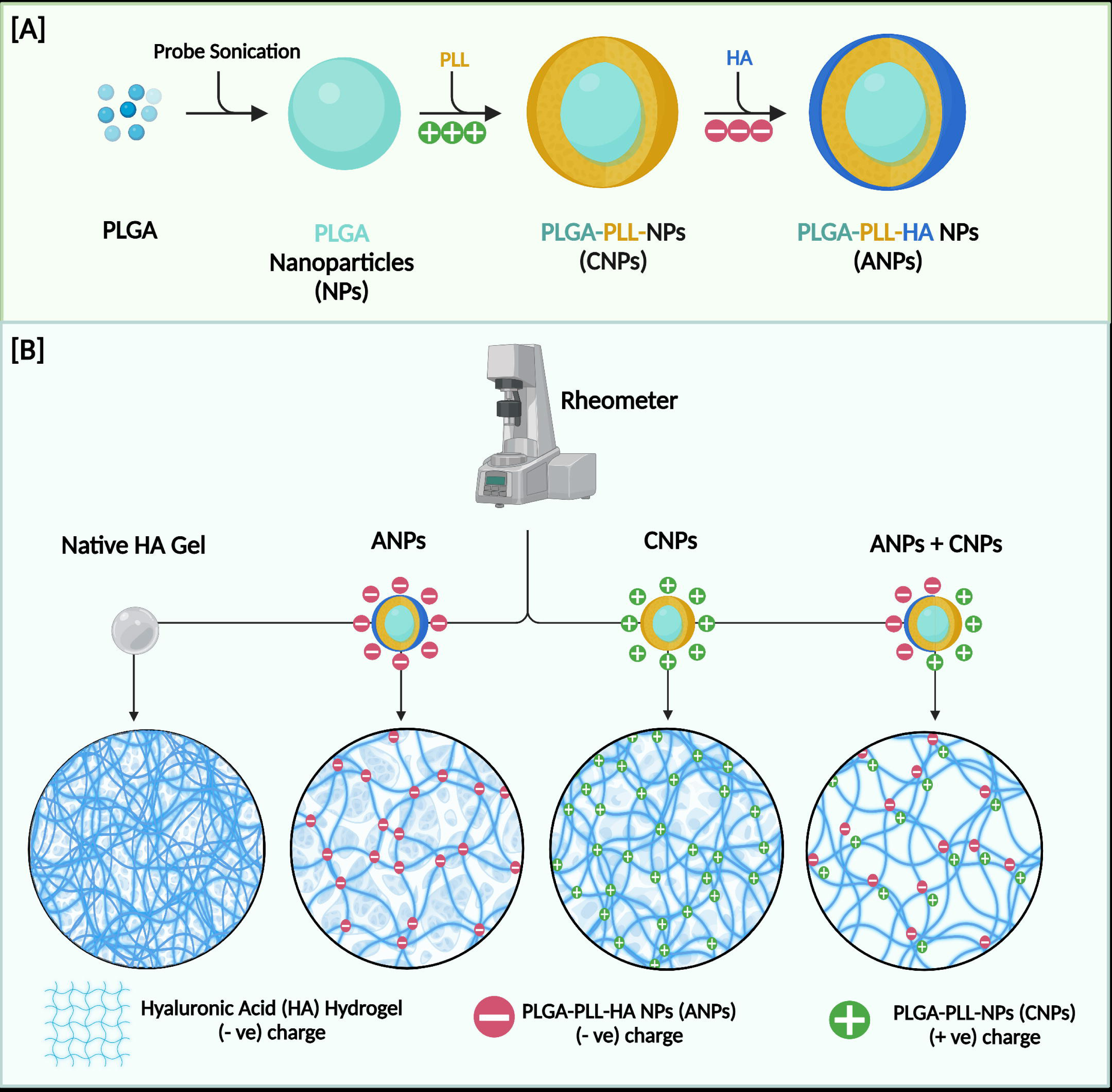
Schematic representation of (A) The fabrication process of the hyaluronic acid (HA) functionalized nanoparticle system, (B) Interaction between the HA gel and the surface charge of the nanoparticle system.

### 2. 3. Fabrication of HA gel-particle system

The hyaluronic acid gel (HA Gel)-particle suspension system was fabricated using HMW HA (1250-1500 k Da). In short, HA (7 mg/mL) was dissolved in PBS (pH 7.4) to obtain native HA gel solution (0.7% w/v) was prepared **(Scheme 1(B))** [1]. Further for each run in DOE experiments, 400 µL of prepared native HA gel solution was used to make particle suspension with CNPs, ANPs and mix of CNPs and ANPs as per the **table 1**.

**Table 1.** The independent variables and their levels as used in the CCD experiments.

| **DOE**<br>**Exp.**<br>**No.** | **Independent**<br>**variables** | **Levels** |  |  | **Star Point $\alpha = 1.41$** |  |
| --- | --- | --- | --- | --- | --- | --- |
|  |  | **Low (-1)** | **Central (0)** | **High (+1)** | **-α** | **+$\alpha$** |
| **1.** | A: ANPs ($\mu\text{L}$) | 20 | 40 | 60 | 11.71 | 68.28 |
|  | B: Amplitude (%) | 4 | 7 | 10 | 2.75 | 11.24 |
| **2.** | C: CNPs ($\mu\text{L}$) | 20 | 40 | 60 | 11.71 | 68.28 |
|  | D: Amplitude (%) | 4 | 7 | 10 | 2.75 | 11.24 |
|  |  | **Levels** |  |  | **Star Point $\alpha = 1.63$** |  |
|  |  | **Low (-1)** | **Central (0)** | **High (+1)** | **-α** | **+$\alpha$** |
| **3.** | E: ANPs ($\mu\text{L}$) | 20 | 40 | 60 | 7.34 | 72.66 |
| | F: CNPs ($\mu\text{L}$) | 20 | 40 | 60 | 7.34 | 72.66 |
|  | G: Amplitude (%) | 4 | 7 | 10 | 2.10 | 11.89 |

### 2. 4. Physicochemical, morphological and rheological analysis of the HA Gel-Particle system

Dynamic light scattering (DLS) using a Lite Sizer 500 (Anton Paar) was employed to measure particle size, polydispersity index (PDI), and surface zeta potential. Microstructural observations of the particles and the HA Gel-particle system were performed using scanning electron microscopy (SEM) with a Hitachi S-4700 EDX. Rheological characterisation of both the native HA gel and the HA gel-particle systems was conducted using an Anton Paar Modular Compact Rheometer (MCR) 302, equipped with a 25 mm parallel plate geometry.

### 2. 5. Experimental design and statistical analysis

The interaction of the surface charge of the nanoparticle with the HA Gel was evaluated using the Design Expert 13 software [20] [21]. The experimental design and analysis were performed using the RSM-CCD. Initially, two experiments with two independent variables were conducted. In the first design experiment, ANPs, and amplitude were selected to get a final response of complex viscosity. Similarly, in the second design experiment, CNPs were used in the place of ANPs. Finaly, to understand the synergistic effect of the interaction of NPs with the hydrogel system, a third set of design experiments was conducted. This set involved three independent variables: ANPs, CNPs, and amplitude, to obtain the eventual response of complex viscosity.

In the first and second set of design experiments, both coded and uncoded values of the two variables are presented in **Table 1**. It is worth emphasising that the interconversion of coded and real values was achieved through the equation below:

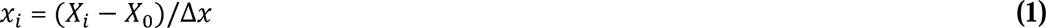

Within the confines of this context, x_i_ serves to represent the dimensionless coded value of the i-th independent variable, X_i_ is used to refer to the actual value of the independent variable, X_0_ designates the actual value of the central point, and Δx denotes the step change value of the variable.

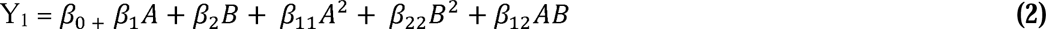

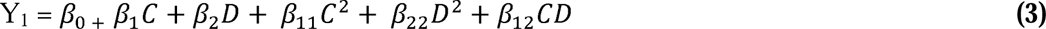

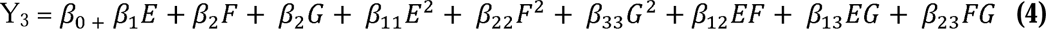

Y_1_, Y_2_, and Y_3_ represent the predicted complex viscosity, while β0 denotes the interactional coefficient. The equation is characterised by the estimated coefficients β1, β2, β3, β11, β22, β33, β12, β13, and β23. The manipulated factors A, B, C, D, E, F, and G correspond to the respective factors provided in the **table 1**.

## 3. Results and discussion

### 3. 1. Physicochemical analysis of the particle system

The differences in size, polydispersity index (PDI), surface zeta potential, particle intensity percentage, and relative frequency percentage of the synthesised polymeric nanoparticle (NPs) system with anionic and cationic coatings were prepared by the layer-by-layer method are depicted in **Fig. 1(A-E)**. The mean sizes of the PLGA NPs, CNPs, and ANPs were 169 ± 12.54 nm, 267 ± 17.87 nm, and 364 ± 41.72 nm, respectively. The initial surface charge of the nanoparticles after sonication was -24 ± 1.70 mV. However, after the PLL coating, the zeta potential significantly shifted to 36 ± 1.18 mV. Upon further coating with HMW-HA, the surface charge changed to -31 ± 1.42 mV.

**Figure 1.**
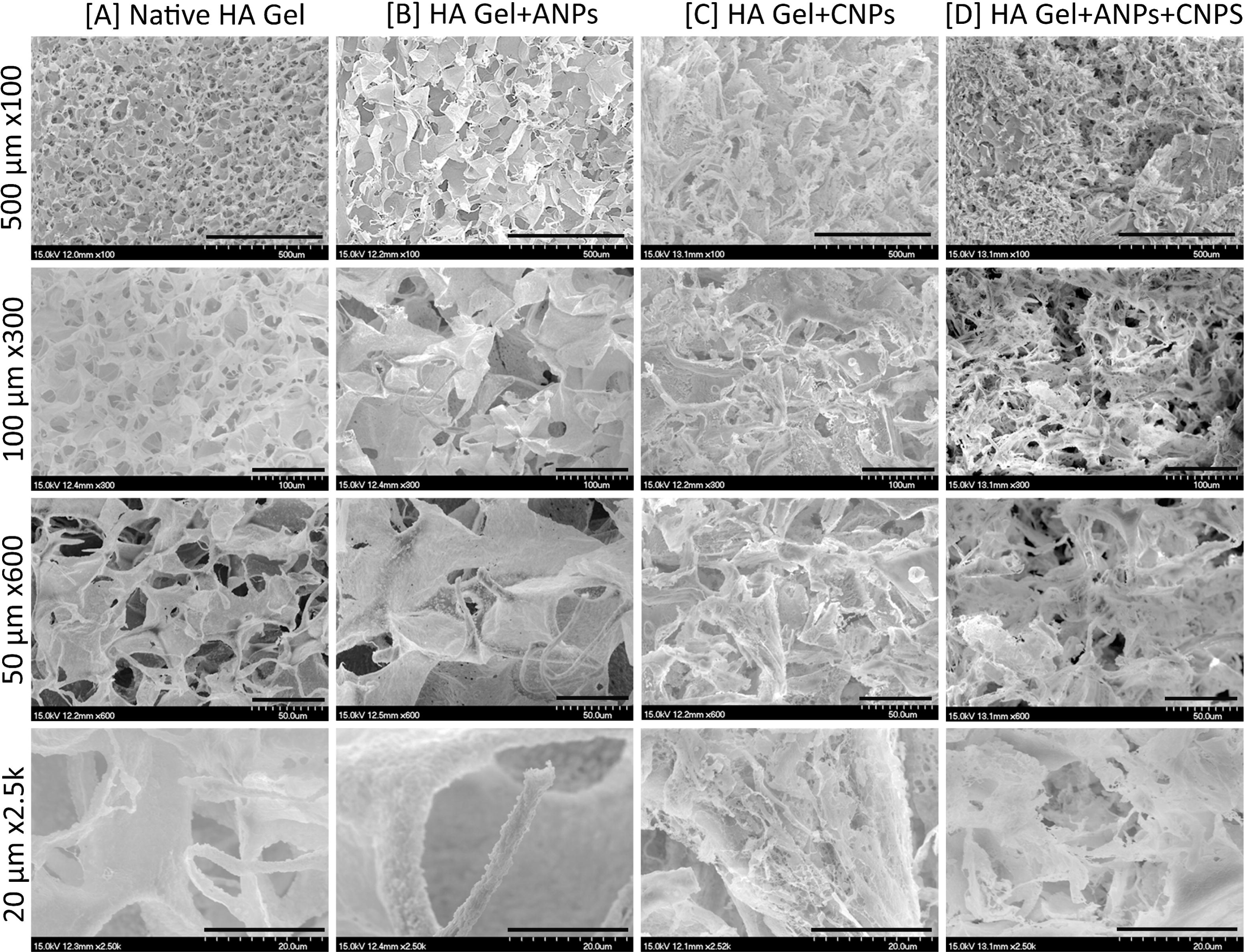
Characterisation of nanoparticle and HA Gel particle system: The difference in (A) Size, (B) Polydispersity (PDI), (C) Surface zeta potential, and the dynamic light scattering (D) intensity and (E) relative frequency % graph of the PLGA NPs, CNPs and ANPs after layer-by-layer coating (data are presented as mean ± S.D (n=3)), (F) Macroscopic images of the HA Gel particle system (G) SEM morphology of the (a) PLGA NPs, (b) CNPs, and (c) ANPs after layer-by-layer coating

### 3. 2. Morphological analysis of the prepared nanoparticle system

The shape and extended morphology of the NPs analysed by SEM was depicted in **Fig. 1G (a-c)**. The prepared NPs (PLGA NPs, CNPs, and ANPs) with different coating exhibited spherical morphology with uniform particle distribution. Significant morphological changes were observed in the SEM images, showing an increase in the size after each layer of coating with a well-defined core-shell structure [22] [23] [24]. The SEM morphology revealed that the NPs were spherical, monodisperse, and ranged in size from 170 to 500 nm, as confirmed by the Lite sizer measurements.

### 3. 3. Highly Interconnected Morphology of the HA Gel-Particle System

The highly interconnected morphology of the HA Gel-particle suspension system with CNPs, ANPs and Mix of CNPs + ANPs was compared with the native HA Gel without particles, as depicted in the macroscopic images in **Fig. 1F**. The SEM micrograph of the HA Gel systems is shown in **Fig. 2(A-D).** The individual and HA Gel-particle system with ANPs **(Fig. 2(C))** exhibited with wider and loosely packed pores with larger void pores sizes compared to the narrower and tightly packed pores with smaller void sizes observed in the HA Gel-particle system with CNPs and mix NPs system **(Fig. 2B and D).** This difference is attributed to the surface charge of the NPs affecting the hydrogel morphology [25]. **Fig. 2A** illustrates the three-dimensional HA Gel-particle system, where particles are seen adhering on the wall of the gel network forming intermolecular hydrogen bonds. This results in a rough surface on the pore walls, in contrast to the smooth and even surface of the native HA Gel [26] [27]. The effect of the NPs on the gel network was related to the hydrogen bonding and electrostatic interaction between the surface charge of the NPs and the HA gel system [28] [29].

**Figure 2.**
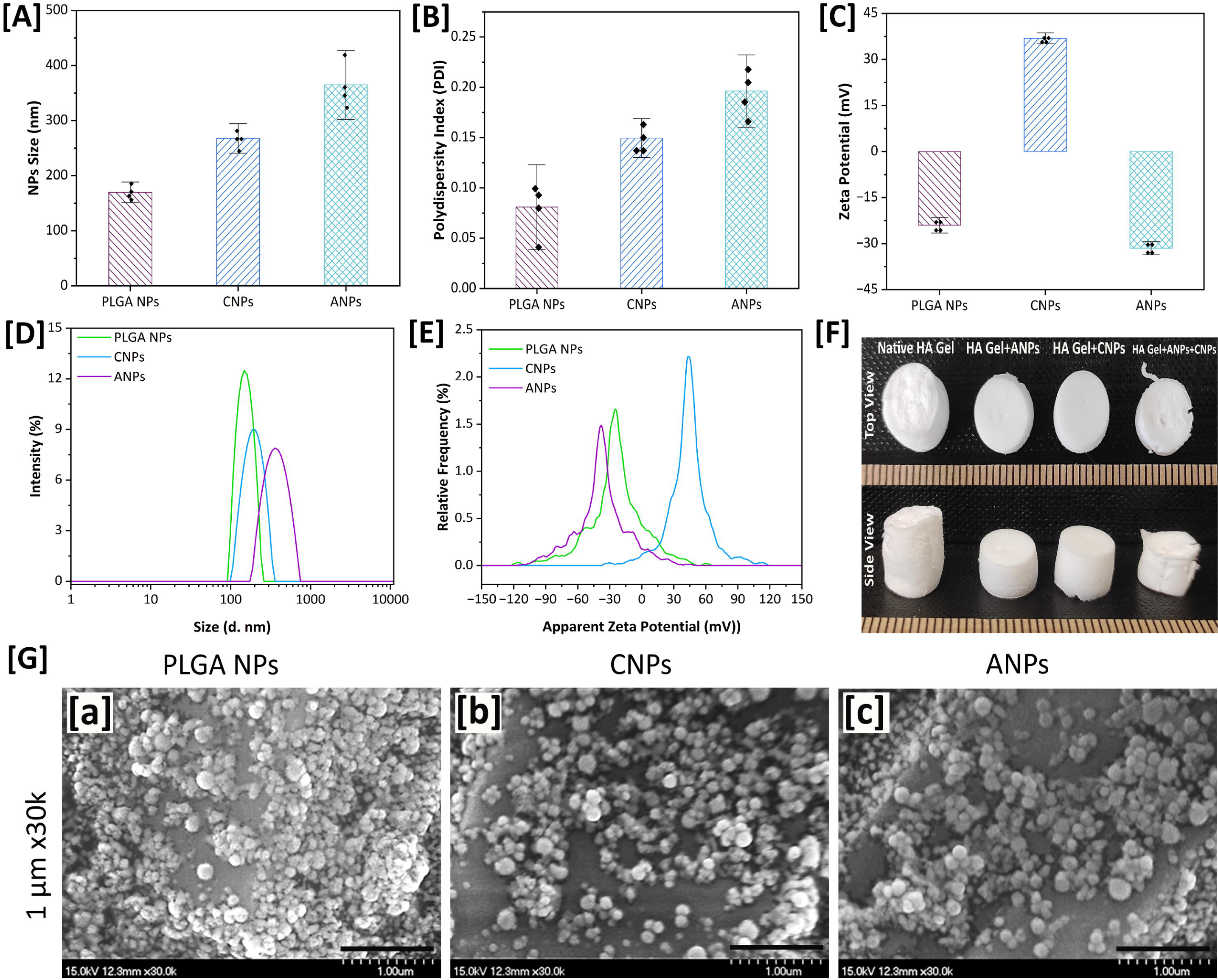
SEM micrographs of the HA Gel system to visualise the microstructural changes under different conditions: (A) native HA Gel, (B) HA Gel with ANPs, (C) HA Gel with CNPs, and (D) HA Gel with ANPs and CNPs at the different magnification (x100, x300, x600, and x2.5k)

### 3. 4. Central composite design

Complex viscosity (η∗) in oscillatory rheology is influenced by various factors, including the volume fraction of nanoparticles and applied shear amplitude. Using Response Surface Methodology (RSM) with a Central Composite Design (CCD), we aimed to optimise the complex viscosity of the HA Gel system through these interactions.

The equation in terms of the actual factors can make predictions about the responses at different levels of each factor in each experiment. In this study, Equation (5) to (7) provided the final equation in terms of the actual factors. Equation (5) shows the mathematical model for predicting the complex viscosity of hydrogel, where the interaction of ANPs with the HA Gel network. Equation (6) depicts the mathematical model for predicting the complex viscosity of hydrogel, where the interaction of CNPs with the HA hydrogel network. The synergistic effect of the interaction between the ANPs and CNPs in the HA hydrogel system was depicted in equation (7) with the mathematical model for predicting the complex viscosity of the hydrogel. In these equations, the positive and negative signs of the coefficients show the positive and negative effects of the obtained response. In all the design experiments, the second-order polynomial model was used to determine the dependent variable (response) using Design Expert software. The model’s validity and adequacy were evaluated through ANOVA to determine significant effects and probable interactions between the variables [20][14].

The ANOVA results of the DOE experiment 1 and 2 were outlined in the **Table S2 and S3**. Based on the ANOVA result from **Table 2**, the interaction of ANPs and CNPs with hydrogel system yielded F-value of 72.28 and 46.05 respectively. Both models were significant, with a p-value of <0.0001. **Table 1** exhibits the ANOVA results from DOE experiment 3, with the F-value of 55. 62, showing the model’s significance. The probability of obtaining F-value of this magnitude because of noise is only 0.01% [20]. A p-value below <0.0001 implies the model significance.

**Table 2.** Analysis of variance (ANOVA) of the response surface quadratic model for HA Gel + ANPs + CNPs system.

| **DOE Exp. No 3** |  |  |  |  |  |
| --- | --- | --- | --- | --- | --- |
| **Source** | **df** | **Sum of Squares** | **Mean Square** | **F-value** | **p-value** |
| **Model** | 9 | 1.134E+05 | 12603.38 | 55.62 | <0.0001 |
| **E** | 1 | 25.65 | 25.65 | 0.1132 | 0.7452 |
| **F** | 1 | 667.41 | 667.41 | 2.95 | 0.1245 |
| **G** | 1 | 2.43 | 2.43 | 0.0107 | 0.9201 |
| **EF** | 1 | 2905.13 | 2905.13 | 12.82 | 0.0072 |
| **EG** | 1 | 46322.07 | 46322.07 | 204.43 | <0.0001 |
| **FG** | 1 | 22.01 | 22.01 | 0.0971 | 0.7633 |
| **E<sup>2</sup>** | 1 | 2835.14 | 2835.14 | 12.51 | 0.0077 |
| **F<sup>2</sup>** | 1 | 36229.60 | 36229.60 | 159.89 | <0.0001 |
| **G<sup>2</sup>** | 1 | 30874.50 | 30874.50 | 136.25 | <0.0001 |
| **Residual** | 8 | 1812.75 | 226.59 |  |  |
| **Lack of Fit** | 5 | 1476.86 | 295.37 | 2.64 | 0.2272 |
| **Pure Error** | 3 | 335.89 | 111.96 |  |  |
| **Cor Total** | 19 | 1.161E+05 |  |  |  |

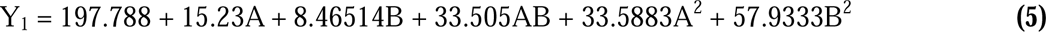

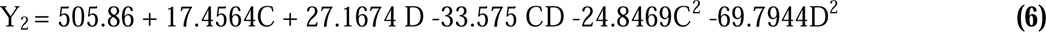

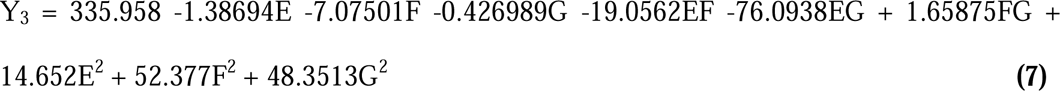

The parameter impact on the response can be determined with the extent from the F-value. Higher F-value shows that the corresponding factors affect the response noticeably [30]. According to the F-value obtained in **table 2**.

The synergistic effect on the volume of the ANPs and CNPs added, possess a significant effect on the complex viscosity when compared to the effect of the NPs individually. Nonetheless, p-value of the lack of fit is not significant, which implies the developed model is more suitable to obtain the complex viscosity by adjusting the volume of NPs with right amplitude for the DOE Exp 1 and 3. However, the DOE exp 2 the lack of fit is significant that implies that model is less suitable to obtained the complex viscosity of desired range because of the increase in the electrostatic attraction between the NPs and the hydrogel network. The ANPs and synergistic effect the NPs possess the strong electrostatic repulsion within the system to obtain the desired complex viscosity [29].

The correlation between the predicted values by the model and the actual results were well correlated with actual complex viscosity obtained. The predicted values are not significantly different from the experimental result suggest that the model with predicted R^2^, standing at 0.8271 alias well, with the adjusted R^2^ of 0.9666. This suggests the reinforcement of the model’s predictability and acceptability. **Fig. 3 (G-I)** shows the model accuracy compares to the experimental results to the mathematically statistically predicted value of the models for obtaining the complex viscosity of the hydrogel as response after the synergistic effect of NPs. The final complex viscosity of the hydrogel was obtained using the 40 µL of both ANPs and CNPs with an amplitude of 7 % as the final parameter to gain the ideal viscosity of the hydrogel with NPs.

**Figure 3.**
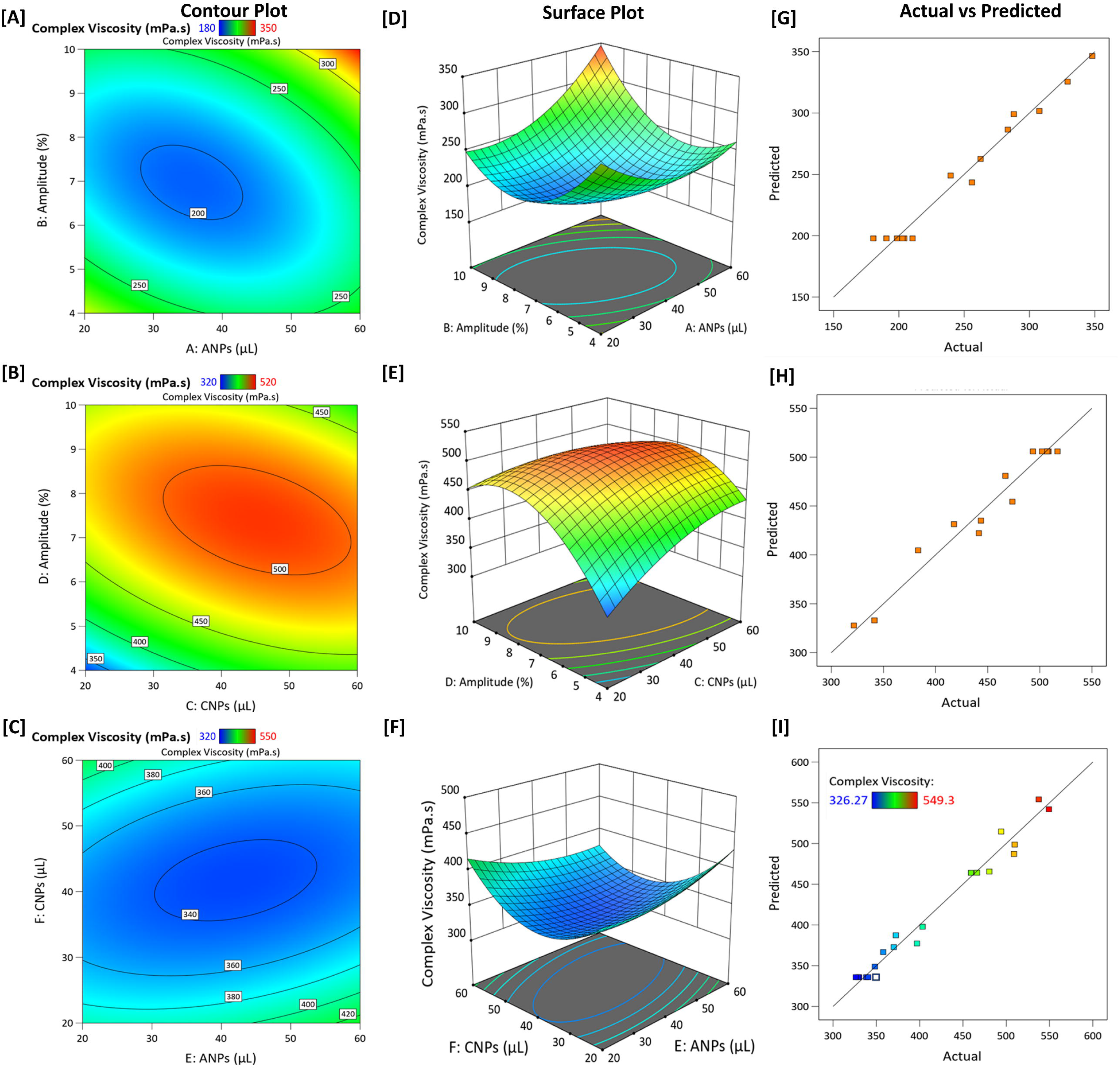
Visualisation of the interactions and effects of variables on complex viscosity in the HA Gel system: (A-C) Contour plots illustrating the relationships and effects of: (A) Volume of ANPs (µL) and amplitude (shear strain %), (B) Volume of CNPs (µL) and amplitude (shear strain %), (C) Volume of ANPs (µL) and Volume of CNPs (µL), (D-F) Surface plots depicting the interaction and effects of: (D) Volume of ANPs (µL) and amplitude (shear strain %) (E) Volume of CNPs (µL) and amplitude (shear strain %), (F) Volume of ANPs (µL) and Volume of CNPs (µL), The complex viscosity (mPa·s) is represented by a color gradient in both the contour and surface plots. The predicted response values for complex viscosity are compared with actual response values for the HA Gel system: (G) Predicted vs. actual response for complex viscosity with ANPs, (H) Predicted vs. actual response for complex viscosity with CNPs, (I) Predicted vs. actual response for complex viscosity with a combination of ANPs and CNPs.

### 3. 4. Effect of the variables

The whole relationship between the independent variables A, B, C, D, E, F, and G with the response variable Y_1_, Y_2_ and Y_3_ can be clearly explained by contour plot and surface plot generated from the empirical predicted equation 5, 6 and 7. The two dimensional (2D) elliptical contour plot (**(Fig. 3(A-C))** depicts the pair-wise combination of two variables by holding the other variable at the centre point level to achieve the mutual interaction of the independent factors. As exhibited from **(Fig. 3(D-F))**, the several surface plots showing the interaction between the targeted variables on complex viscosity by changing them into 3D diagram. These interactions were developed to achieve the optimum condition to obtain the desired complex viscosity. Hence, in all plots, fixed variables are held at ANPs (40 µL), CNPs (40 µL) and amplitude (7%). **Fig. 3 (A and D)** shows the interactive effects of ANPs, and amplitude gives with decreased complex viscosity.

The presence of negatively charged surfaces on the NPs leads to significant electrostatic repulsion with the negatively charged HA hydrogel, resulting in a decrease in complex viscosity because of the disruption of the hydrogel network structure [1] As shown in **Fig. 3 (B and E)**, introducing CNPs and the variation in amplitude increase the complex viscosity of the hydrogel. This is attributed to the strong electrostatic attraction between the positively charged CNPs and the negatively charged HA hydrogel, which enhances the network structure and thus increases the complex viscosity [17]. CNPs on the hydrogel are significantly high than the effect of ANPs on the hydrogel. Furthermore, **fig. 3 (C and F)** clearly depict the synergist effect of both ANPs and CNPs on the hydrogel, that clearly depicted as contour and surface plot. The negative charge on the surface of the NPs dominated over the positive charge on the NPs to achieve a considerably complex viscosity in between the range of complex viscosity achieved by individual NPs effect. Tiffany et al. reported that the viscosity of a material increases when the surface charge of particles transitions from anionic to cationic in a particle gel system [31]. Additionally, the interaction of chitosan nanoparticles with the HA hydrogel demonstrated significant potential, resulting in a stiffer and more rigid structure compared to pure HA gel [18]. Similar effects of ANPs and CNPs with varying amplitude are shown in **Fig. S1**, further supporting the observed trends.

### 3. 5. Rheological properties HA Gel-Particle system

#### 3. 5. 1. Preliminary Characterisation of HA Gel

Preliminary experiments involving amplitude (1 to 20% shear strain) and frequency sweeps (100 to 0.1 rad/s) were conducted to determine the linear viscoelastic region and deformation behaviour of the native HA gel at 37 °C. **Fig. 4(A)** illustrates the deformation behaviour of the HA gel with increasing applied amplitude. **Fig. 4(B)** demonstrates the transition of native HA gel from elastic (G’ > G’’) to viscous (G’ < G’’) behaviour. The crossover point, where G’ equals G’’, signifies this transition and provides insights into the relaxation properties of the native HA gel. Following this initial characterisation, a time sweep experiment was conducted at a constant amplitude and frequency to evaluate the complex viscosity of the native HA gel. The HA gel concentration (0.7%) and angular frequency (1 rad/s) were kept constant throughout these experiments.

**Figure 4.**
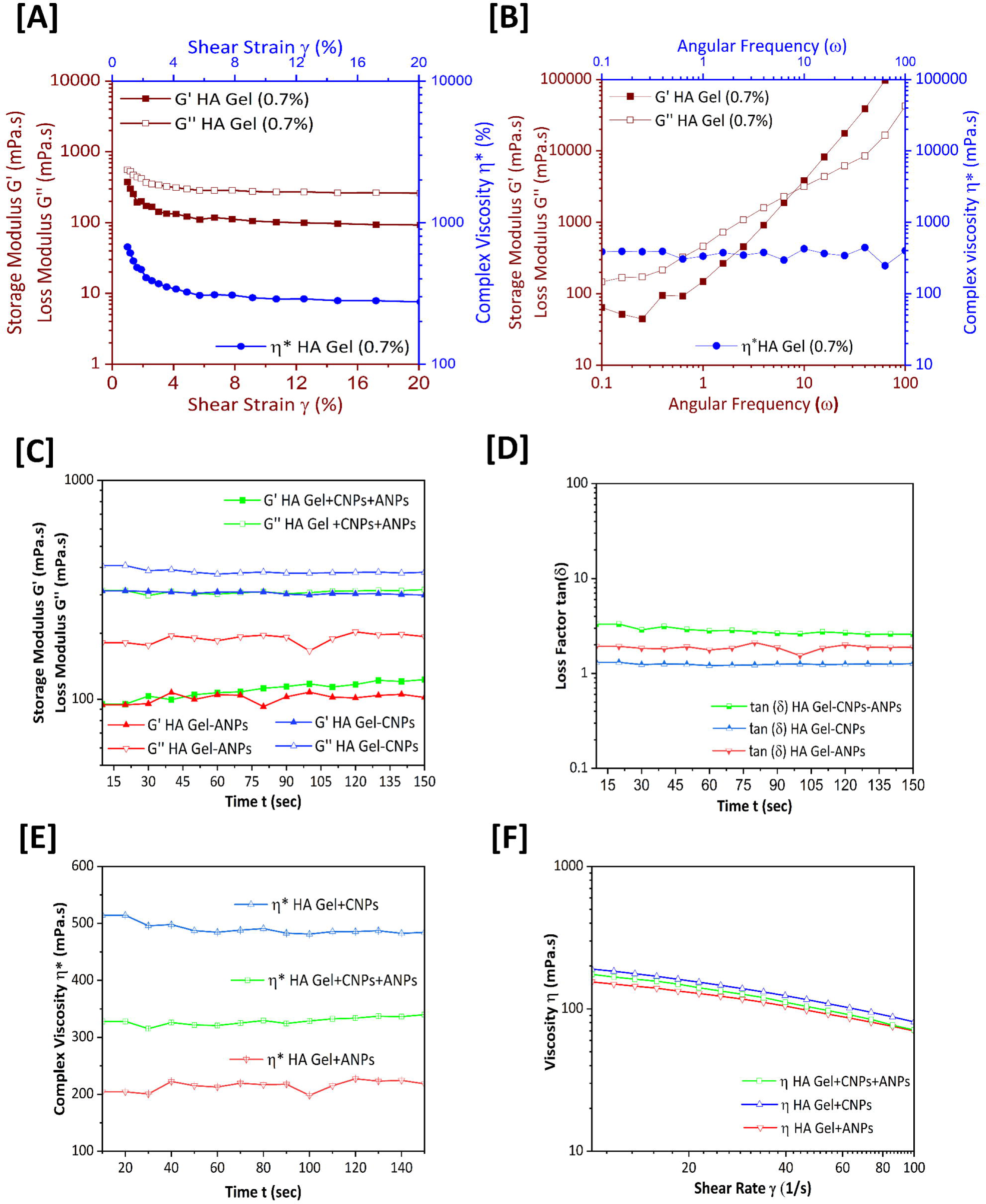
Characterisation of the native HA gel system showing rheological properties: Storage modulus (G’) and loss modulus (G’) at (A) constant angular frequency (rad/s), (B) constant amplitude (shear stress, %), (C) Comparative analysis of storage modulus (G’) and loss modulus (G’) across at a constant amplitude of 7% and an angular frequency of 1 rad/s over time (seconds), (D) Loss modulus represented as tan δ, (E) Complex viscosity (η*) measured at a constant amplitude of 7% and an angular frequency of 1 rad/s over time (seconds), (F) Viscosity (η) as a function of shear rate, ranging from 10 to 100 s[¹, for the HA Gel Particle system (HA Gel + ANPs, HA Gel + CNPs, and HA Gel + ANPs + CNPs). The data represents the mean viscosity (± S.D.) for n=3 samples.

#### 3. 5. 2. Complex Viscosity behaviour of the HA Gel-Particle system

In the experimental design, the volume of CNPs and ANPs, along with amplitude, was systematically varied to investigate their impact on the resulting complex viscosity. The viscoelastic behaviour of the HA gel-particle system revealed that G’’ > G’, indicating that the interaction between the particles and the gel did not decimate the complex viscosity. A significant decrease in complex viscosity was observed in the HA gel with ANPs, attributed to the disruption of the HA gel polymeric network by strong electrostatic repulsion between the particles and the gel system. This observation is consistent with Kotla et al., who noted a decrease in complex viscosity because of the breakdown of the hydrogel polymeric network when anionic nanoparticles were introduced [1] [32]. Conversely, the HA gel with CNPs exhibited a significant increase in complex viscosity compared to the native HA gel, likely because of the electrostatic attraction between the polymeric network and CNPs **(Fig. 4(C, D, and E))** [18]. The synergistic effect of ANPs and CNPs in the HA gel system resulted in a mixed response, with the resulting complex viscosity falling within the range observed for the individual CNP and ANP systems. This suggests an interplay between attractive and repulsive forces in the mixed NP system, ultimately influencing the complex viscosity of the HA gel [29].

#### 3. 5. 3. Viscosity behaviour of the HA Gel-Particle system

The individual and synergistic effects of the HA gel-particle system exhibited a consistent trend in both viscosity and complex viscosity compared to the native HA gel, as shown in **Fig. 4(F)**. The viscosity decreased with increasing shear rate, showing that the HA gel with particles exhibits shear thinning behaviour (non-Newtonian) [33] [34]. The dominant attractive forces between particles increase the viscosity, while dominant repulsive forces decrease it. The volume and size of the particles also affect the material’s viscosity. The surface charge of the particles decreases when interacting with the HA gel system, leading to a loss of monodisperse suspension and the formation of aggregates [35] [36].

## 4. Conclusion

The optimisation of the complex viscosity of the HA Gel-particle system was successfully achieved using response surface methodology within a central composite design framework. This study highlights that the surface charge of nanoparticles significantly influences the complex viscosity of the HA Gel system. Specifically, anionic nanoparticles decrease the viscosity, while cationic nanoparticles increase it. The mixture of both types of nanoparticles results in an intermediate viscosity. Through meticulous analysis of the interactions between nanoparticles and the HA Gel matrix, it was discovered that the volume fraction and amplitude of nanoparticles are essential rheometric process parameters that directly impact complex viscosity. These findings pave the way for advanced biomedical pharmaceutical formulations, enabling precise control over gel properties for rectal gel enema applications. This approach provides a robust method for tailoring the viscoelastic properties of hydrogel systems, enhancing their functionality and efficacy in targeted medical applications.

## Acknowledgements

The authors would like to thank Centre for Microscopy and Imaging, university of Galway. This work was supported by a grant from the from Science Foundation Ireland (SFI) and the European Regional Development Fund (ERDF) under Grant number 13/RC/2073_P2.

## CRediT authorship contribution statement

**Giriprasath Ramanathan:** Conceptualisation, Methodology, Investigation, Validation, Data curation, Writing - Original Draft, Writing – review & editing, Visualisation. **Masroora Hassan:** Conceptualisation, Investigation, Formal analysis, Writing - Review & Editing. **Yury Rochev:** Conceptualisation, Writing – review & editing, Resources, Supervision, Validation, Funding acquisition, Project administration.

