## Supplementary Material for "Optimising the viscoelastic properties of hyaluronic acid hydrogels through colloidal particle interactions: a response surface methodology approach"

**Electronic supplementary material**

**Table S1. The design of experiments used in this study with their experimental response for complex viscosity (CV) by the CCD method**

| DOE Exp. No 1. | | | | DOE Exp. No 2. | | | | DOE Exp. No 3. | | | | |
|---|---|---|---|---|---|---|---|---|---|---|---|---|
| Run order | Independent variables | | Response | Run order | Independent variables | | Response | Run order | Independent variables | | | Response |
| | A: ANPs (μL) | B: A (%) | $Y_1$: CV (mPa.s) | | C: CNPs (μL) | D: A (%) | $Y_2$: CV (mPa.s) | | E: ANPs (μL) | F: CNPs (μL) | G: A (%) | $Y_3$: CV (mPa.s) |
| **1** | 40 | 7 | 210.44 | **1** | 40 | 7 | 502.15 | **1** | 40 | 40 | 7 | 349.57 |
| **2** | 60 | 4 | 262.52 | **2** | 60 | 4 | 443.51 | **2** | 60 | 20 | 10 | 403.39 |
| **3** | 20 | 10 | 239.59 | **3** | 20 | 10 | 473.72 | **3** | 40 | 40 | 7 | 330.09 |
| **4** | 20 | 4 | 288.02 | **4** | 20 | 4 | 341.43 | **4** | 60 | 60 | 4 | 509.81 |
| **5** | 60 | 10 | 348.11 | **5** | 60 | 10 | 441.50 | **5** | 20 | 20 | 4 | 357.94 |
| **6** | 40 | 7 | 180.42 | **6** | 40 | 7 | 508.28 | **6** | 20 | 60 | 10 | 549.30 |
| **7** | 40 | 7 | 190.42 | **7** | 40 | 7 | 493.41 | **7** | 60 | 60 | 10 | 348.34 |
| **8** | 40 | 11.24 | 329.21 | **8** | 40 | 11.24 | 383.23 | **8** | 20 | 20 | 10 | 494.21 |
| **9** | 40 | 7 | 202.75 | **9** | 40 | 7 | 516.96 | **9** | 40 | 40 | 7 | 329.03 |
| **10** | 40 | 7 | 204.19 | **10** | 40 | 7 | 507.03 | **10** | 20 | 60 | 4 | 372.48 |
| **11** | 11.71 | 7 | 255.99 | **11** | 11.71 | 7 | 417.67 | **11** | 40 | 40 | 7 | 338.34 |
| **12** | 40 | 7 | 198.51 | **12** | 40 | 7 | 507.06 | **12** | 60 | 20 | 4 | 537.58 |
| **13** | 68.28 | 7 | 283.44 | **13** | 68.28 | 7 | 467.02 | **13** | 40 | 40 | 7 | 326.27 |
| **14** | 40 | 2.75 | 307.60 | **14** | 40 | 2.75 | 321.67 | **14** | 40 | 40 | 7 | 340.61 |
| | | | | | | | | **15** | 72.65 | 40 | 7 | 370.15 |
| | | | | | | | | **16** | 40 | 7.34 | 7 | 508.97 |
| | | | | | | | | **17** | 40 | 72.65 | 7 | 459.28 |
| | | | | | | | | **18** | 40 | 40 | 2.10 | 480.47 |
| | | | | | | | | **19** | 7.34 | 40 | 7 | 396.90 |
| | | | | | | | | **20** | 40 | 40 | 11.89 | 466.31 |

**Table S2. Analysis of variance (ANOVA) of the response surface quadratic model for HA Gel + ANPs system**

| DOE Exp. No 1. | | | | | |
|---|---|---|---|---|---|
| **Source** | **df** | **Sum of Squares** | **Mean Square** | **F-value** | **p-value** |
| **Model** | 5 | 38008.41 | 7601.68 | 72.28 | <0.0001 |
| **A** | 1 | 1855.63 | 1855.63 | 17.64 | 0.0040 |
| **B** | 1 | 573.27 | 573.27 | 5.45 | 0.0523 |
| **AB** | 1 | 4490.34 | 4490.34 | 42.70 | 0.0003 |
| **$A^2$** | 1 | 8331.15 | 8331.15 | 79.21 | <0.0001 |
| **$B^2$** | 1 | 24784.77 | 24784.77 | 235.66 | <0.0001 |
| **Residual** | 7 | 736.20 | 105.17 | | |
| **Lack of Fit** | 3 | 251.43 | 83.81 | 0.6915 | 0.6033 |
| **Pure Error** | 4 | 484.77 | 121.19 | | |
| **Cor Total** | 13 | 39020.69 | | | |

**Table S3. Analysis of variance (ANOVA) of the response surface quadratic model for HA Gel + CNPs system**

| DOE Exp. No 2. | | | | | |
|---|---|---|---|---|---|
| **Source** | **df** | **Sum of Squares** | **Mean Square** | **F-value** | **p-value** |
| **Model** | 5 | 51642.18 | 10328.44 | 46.05 | <0.0001 |
| **C** | 1 | 2437.82 | 2437.82 | 10.87 | 0.0132 |
| **D** | 1 | 5904.53 | 5904.53 | 26.33 | 0.0014 |
| **CD** | 1 | 4509.12 | 4509.12 | 20.11 | 0.0029 |
| **$C^2$** | 1 | 4559.02 | 4559.02 | 20.33 | 0.0028 |
| **$D^2$** | 1 | 35972.34 | 35972.34 | 160.39 | <0.0001 |
| **Residual** | 7 | 1569.93 | 224.28 | | |
| **Lack of Fit** | 3 | 1394.44 | 464.81 | 10.59 | 0.0225 |
| **Pure Error** | 4 | 175.49 | 43.87 | | |
| **Cor Total** | 13 | 53705.25 | | | |

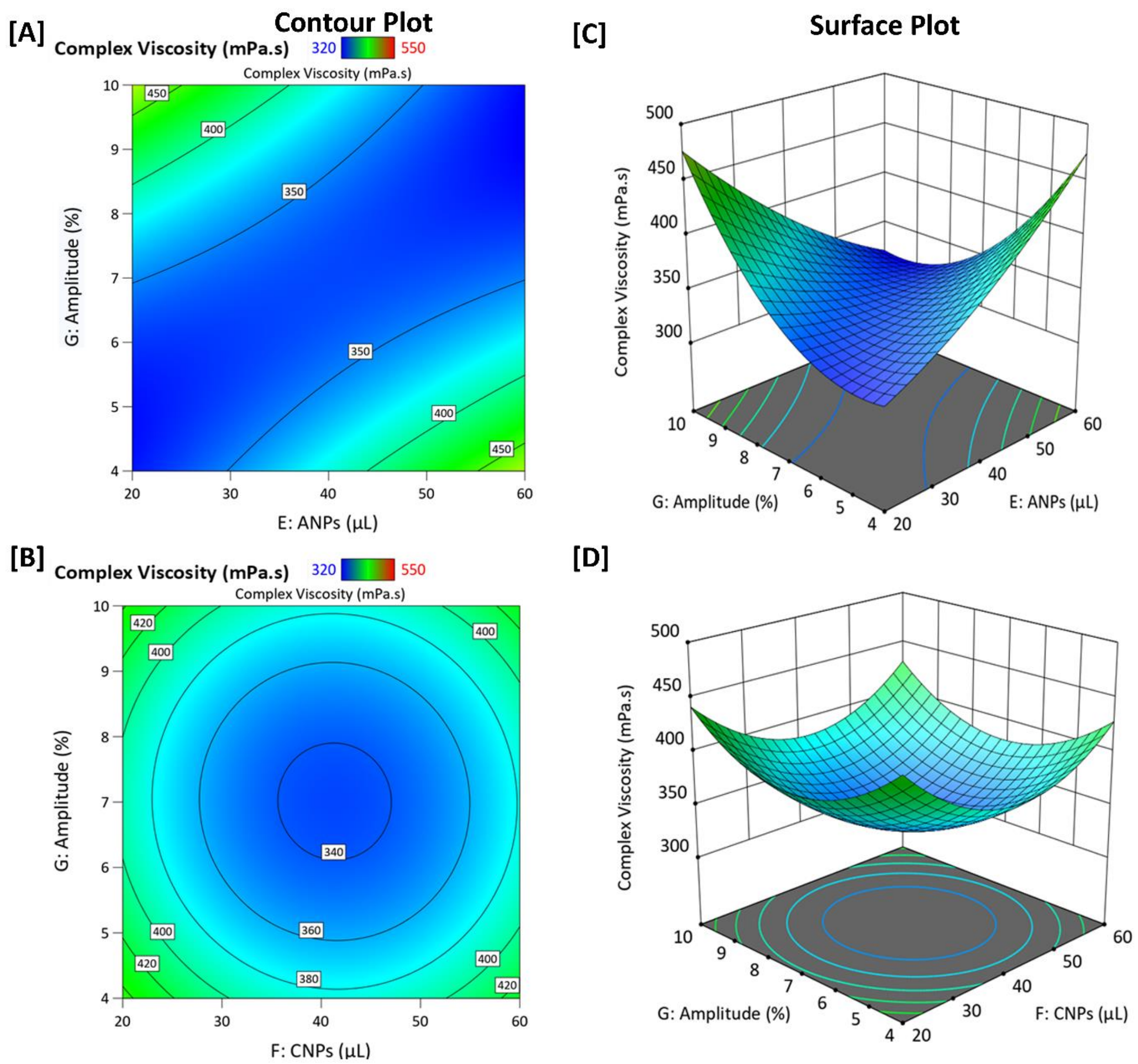


**Figure S1. (A-B) Contour plots and (C-D) surface plots illustrating the interaction and effects of variables on complex viscosity. (EG) Volume of anionic nanoparticles (ANPs, µL) and amplitude (shear strain %); (FG) Volume of cationic nanoparticles (CNPs, µL) and amplitude (shear strain %). The complex viscosity (mPa·s) range is represented by the color gradient.**
